# Strong and tunable anti-CRISPR/Cas9 activity of AcrIIA4 in plants

**DOI:** 10.1101/2021.01.08.425920

**Authors:** Camilo Calvache, Marta Vazquez-Vilar, Sara Selma, Mireia Uranga, José Antonio Daròs, Diego Orzáez

## Abstract

This study describes the strong anti-CRISPR activity of the bacterial AcrIIA4 protein in *Nicotiana benthamiana*, a model plant used as molecular farming platform. The results demonstrate that AcrIIA4 abolishes site-directed mutagenesis in leaves when transiently co-expressed with CRISPR/Cas9. We also show that AcrIIA4 represses CRISPR/dCas9-based transcriptional activation (CRISPRa) of both reporter and endogenous genes in a highly efficient, dose-dependent manner. Furthermore, the fusion of an auxin degron to AcrIIA4 results in auxin-regulated activation of a downstream reporter gene. The strong anti-Cas9 activity of AcrIIA4 reported here opens new possibilities for customized control of gene editing and gene expression in plants.

## Main

The systems based on clustered regularly interspaced short palindromic repeats (CRISPR) and CRISPR associated (Cas) proteins have been used as the site-directed nuclease of choice for plant genome engineering due to its simplicity and specificity^1^. In addition to gene editing through site-specific DNA cleavage and subsequent repair by the host machinery, CRISPR/Cas-based platforms provide new ways of engineering other catalytic and non-catalytic genome-associated functions, such as DNA base modifiers, epigenetic effectors or programmable transcription factors (PTFs); the latter enabling new control strategies for gene expression known as CRISPR activation (CRISPRa) or repression (CRISPRi)^2,3^. This is achieved by coupling nuclease-deactivated versions of Cas (dCas) to transcriptional activator or repressor domains, which then act as functionally-customized RNA-guided DNA-binding complexes^2,4^.

Although different CRISPR-based architectures in plants have been shown to induce both efficient mutagenesis and powerful transcriptional activation when expressed constitutively^5,6^, the spatial-temporal control on such activities would considerably expand the range of their potential applications. To this date, the control of CRISPR/Cas activity in plants has been attempted mainly at the transcriptional level using inducible promoters controlling expression of Cas transcripts^7–9^. However, in practical terms, the control of effector function at the transcriptional level alone is only partially effective due to inherent leakiness and noise. Furthermore, transcriptional regulation imposes a lag in the responses due to the requirement for *de novo* protein synthesis. Conversely, many biological switches make use of post-translational regulatory strategies such as enzymatic modification or protein-protein interactions as relay mechanisms. In this context, the recent discovery of phage-derived proteins with Cas-inhibitory activities, named collectively as anti-CRISPR (Acr) proteins offers exciting new possibilities to control Cas activity at the post-translational level. Acr proteins constitute a collective arsenal of naturally evolved CRISPR/Cas antagonist that inhibit CRISPR/Cas immune activity at various stages^10–12^. To date, 45 non-homologous Acr proteins (24 for class I CRISPR/Cas and 21 for class II) have been discovered, comprising different mechanisms and structures^13–15^. Acr proteins can inhibit CRISPR/Cas function by directly interacting with a Cas protein to prevent DNA binding, crRNA loading, cleavage, or by interfering with effector-complex formation^12^. Whereas Acrs have been successfully tested in mammalian and bacterial cells, no evidence of their functionality in plants exists. Here we study the ability of an Acr protein from a *Listeria monocytogenes* prophage (AcrIIA4) to inhibit the activity of *Streptococcus pyogenes* Cas9 (SpCas9) in *Nicotiana benthamiana*.

First, we analyzed the ability of AcrIIA4 to prevent Cas9 site-specific mutagenesis in plant tissues using three previously well characterized targets: the xylosyltransferase gene (XT) and two different genes of the Squamosa-promoter binding protein-like (SPL) family (Supplementary Table 1). As shown in Fig. 1A and 1B, transient expression of specific gRNAs along with the Cas9 resulted in average editing efficiencies of 6%, 8% and 10% for each target respectively, whereas co-infiltrations of the same editing constructs with a transcriptional unit (TU) expressing a nuclear-localized AcrIIA4 under the constitutive CaMV 35S promoter reduced the editing efficiencies to undetectable levels in all three targets. These results confirmed the ability of AcrIIA4 to prevent Cas9-mediated editing.

**Figure 1.**
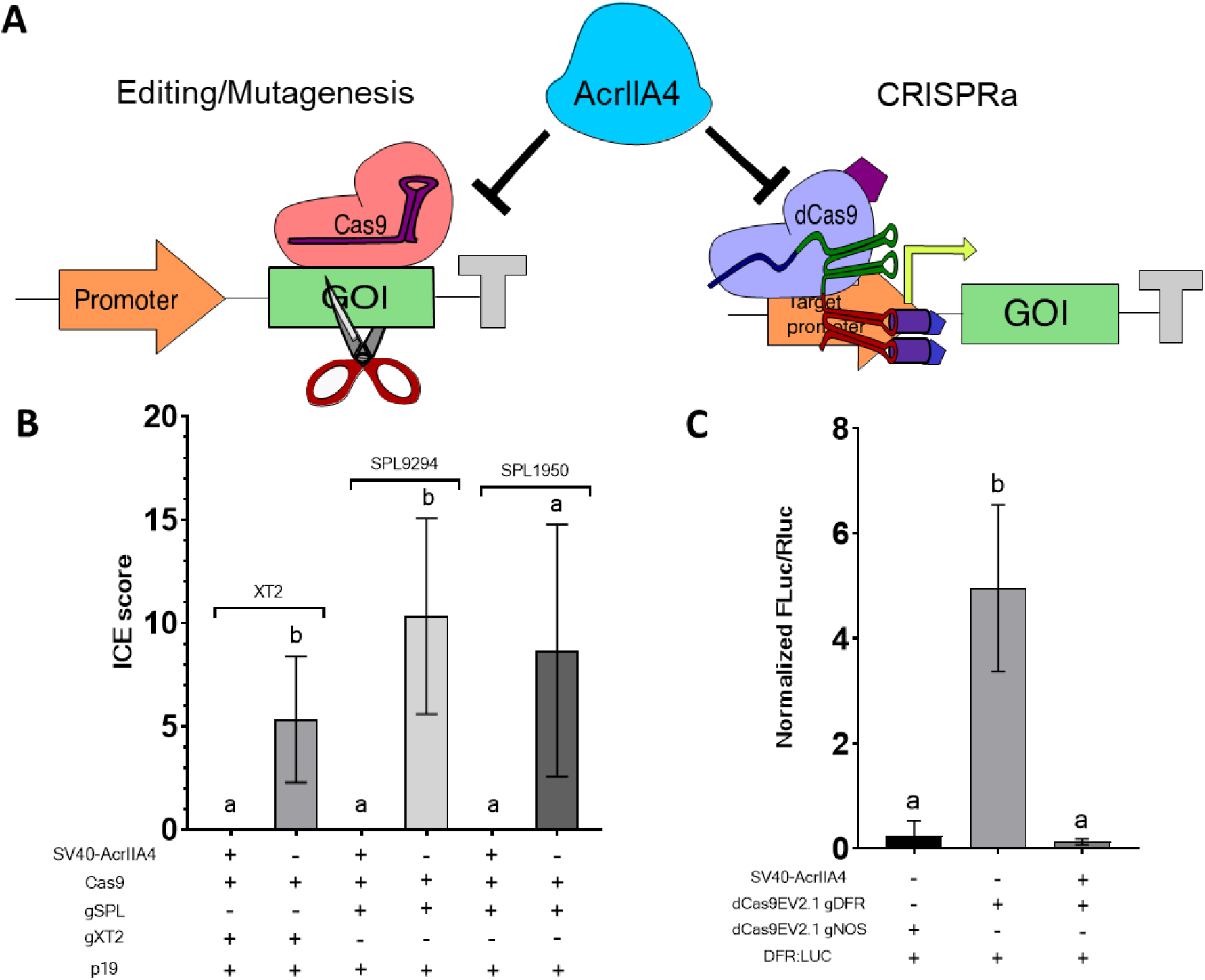
AcrIIA4 prevents Cas9 editing in *N. benthamiana.* A) Schematic representation of AcrIIA4 activity. B) Editing efficiency of *NbXT2* and *NbSPL* in the absence and in the presence of SV40-AcrIIA4. *N. benthamiana* leaves were agroinfiltrated with a Cas9 TU, a P19 TU and the corresponding gRNA, with or without the SV40-AcrIIA4 TU. Editing efficiencies were determined with Synthego. C) CRISPRa repression of SV40-AcrIIA4. Bars show normalized Fluc/Rluc ratios of *N. benthamiana* leaves expressing dCasEV2.1 and the corresponding specific or unspecific gRNAs, with or without AcrIIA4. Error bars indicate SD (n=3). Statistical analyses were performed using unpaired t-Test (P Value <0.05). Variables carrying not significant differences are coupled within the same statistical groups marked with the same letters.

dCas9-based programmable transcriptional activators (PTAs) are becoming widely used in plants for customized regulation of gene expression. We previously developed a collection of dCas9-based PTAs for plant gene control, being the strongest one the so-called dCasEV2.1 activator complex^16^. dCasEV2.1 comprises three elements, each one encoded in a different TU: (i) a modified scaffold RNA, which includes anchoring sites for the phage MS2 coat protein (MCP)^17^; (ii) a dCas9 fused to the EDLL plant activator domain, and (iii) the MCP fused to the synthetic VP64-p65-Rta (VPR) activator domain. We first validated the ability of AcrIIA4 to prevent dCasEV2.1-based gene activation in plants using a transient assay based on a luciferase reporter. The reporter system comprised a Firefly luciferase CDS (Fluc) driven by the *Solanum lycopersicum* DFR promoter (pSlDFR) and a dCasEV2.1 complex targeted to position −147 of the pSlDFR. Luminescence assays showed that the relative transcription activity (RTA) conferred by pSlDFR changed from 0.25 ± 0.28 in its basal state (with dCasEV2.1 loaded with an unrelated gRNA) to 4.96 ± 1.59 in the activated stage (with dCasEV2.1 loaded with a promoter-specific gRNA Fig. 1B). Remarkably, co-expression of AcrIIA4 along with dCasEV2.1 and the pSlDFR-specific gRNA reduced RTA to 0.13 ± 0.26 (Fig. 1A and 1C).

It has been suggested that AcrIIA4 inhibits Cas9 activity by a binding mimicking mechanism, occupying the PAM DNA-binding site with higher affinity than the DNA substrate^18^. Accordingly, its anti-Cas activity is expected to be dependent on AcrIIA4 concentration. To test this hypothesis, we performed a dose-response experiment, infiltrating decreasing amounts of the agrobacterium culture that drives expression of AcrIIA4. As expected, AcrIIA4 anti-Cas9 activity showed a strong dose-dependent effect (Figure 2A). The two highest optical densities (OD_600_) assayed, namely 0.1 and 0.05, resulted in the complete inhibition of the dCasEV2.1-mediated activation, with RTA values of ~0.09. Based on previous estimations of the multiplicity of transformation (MOT) as a function of the agrobacterium concentration in *N. benthamiana* leaf agroinfiltration^19^, we estimated that a complete dCasEV2.1 inhibition was reached with between five and seven T-DNA copies per cell. The lowest OD_600_ that resulted in a significant reduction of the dCasEV2.1-mediated activation was 0.005 (equivalent to an estimated three T-DNA copies per cell on average). This is a bacterial concentration 20 times lower than that used for transferring the dCasEV2.1 construct, which leads to approximately seven T-DNA copies per cell on average. Considering that all transcriptional units in the assay are under the control of the CaMV35S promoter, these results suggest a strong anti-Cas9 activity of AcrIIA4 in plants, a very efficient expression of AcrIIA4, or both.

**Figure 2.**
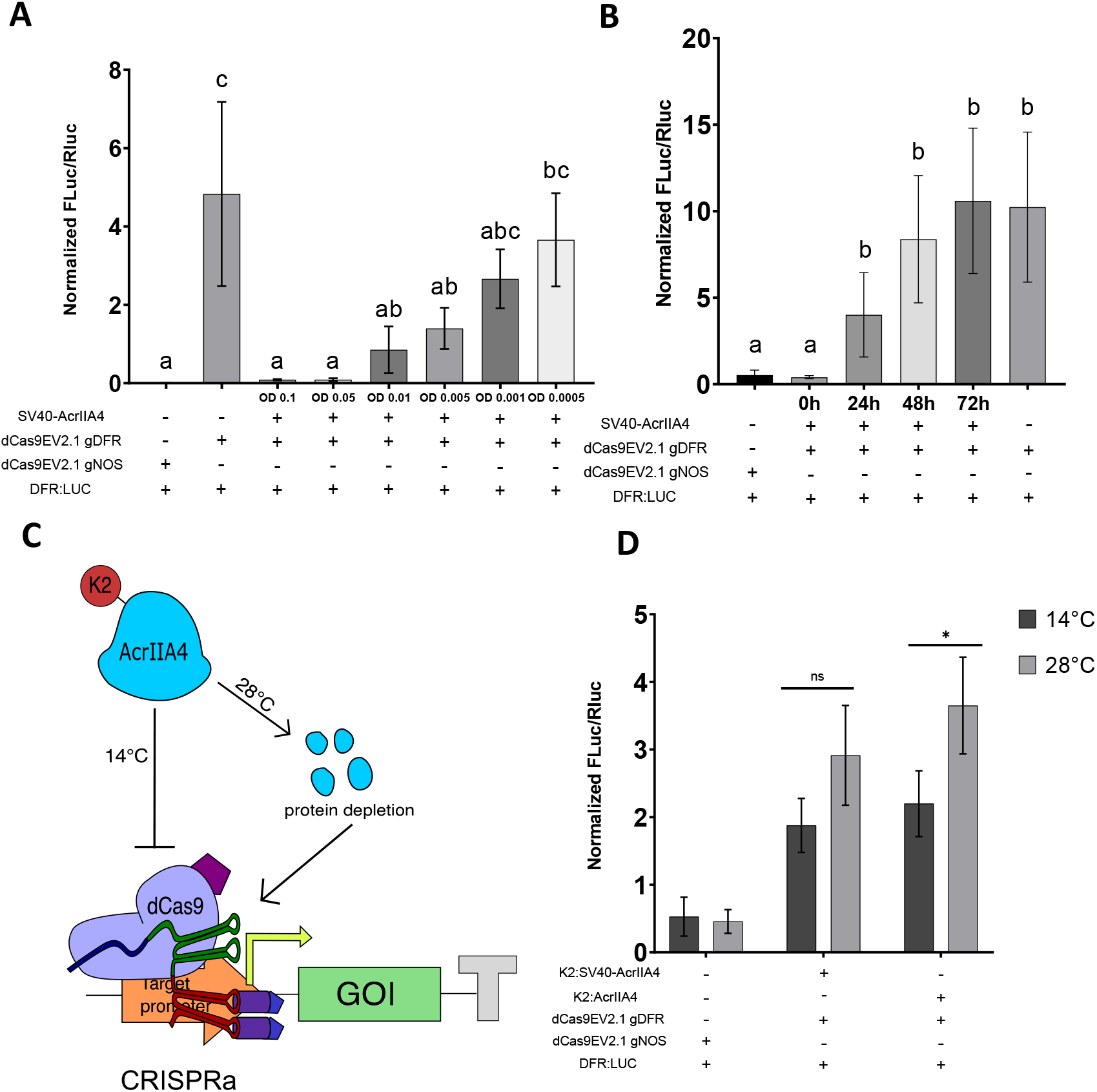
AcrIIA4 prevents dCas9-based activation of a reporter gene in *N. benthamiana* in a regulable manner. A) Normalized Fluc/Rluc ratios of *N. benthamiana* leaves expressing dCasEV2.1 and the corresponding specific or unspecific gRNAs, with AcrIIA4 infiltrated at different OD600 B) Normalized Fluc/Rluc ratios of *N. benthamiana* leaves expressing dCasEV2.1 and the corresponding specific or unspecific gRNAs, with AcrIIA4 infiltrated at different time points. C) Schematic representation of AcrIIA4 at high temperature driven by the K2 low temperature degron. D) Changes in transcriptional de-activation measured as normalized Fluc/Rluc ratios of *N. benthamiana* leaves expressing the low temperature degron K2-AcrII4 fusions in response to low (14°C) and high (28°C) temperature treatments. SV40 is a nuclear localization signal. Error bars indicate SD (n=3). Statistical analyses were performed using One-way ANOVA (Tukey’s multiple comparisons test, P-Value ≤ 0.05,) or unpaired t-Test for two samples comparisons (*P-value ≤ 0.05 and **P-value ≤ 0.005).

A key factor to consider when regulating different components in a genetic circuit is time course dependency. Therefore, we decided to test the ability of the AcrIIA4 to suppress an ongoing dCasEV2.1 activation. For this analysis, AcrIIA4 construct was infiltrated simultaneously, or with a delay of 24, 48 and 72 h after the infiltration of the reporter/dCasEV2.1/gRNA mix, and all tissue samples were subsequently collected 96 h after the initial infiltration. As shown in Fig. 2B, anti-Cas9 treatment is effective up to 24h post initial infiltration, but this efficiency is substantially lost at longer timepoints. These results indicate that displacement of dCas9-DNA activation complexes by AcrIIA4 is a relatively slow or inefficient process and that PTA inhibition by AcrIIA4 is more successfully achieved by early formation of inhibitory complexes rather than by binding competition.

As a first attempt to regulate anti-CRISPR activity post-translationally, we created and assayed an N-terminal fusion of the AcrIIA4 with the temperature-dependent K2 degron, which was earlier described to induce temperature-dependent degradation when translationally fused to other proteins-of-interest^20^ (Fig. 2C). We assayed two K2 configurations with or without the nuclear localization signal SV40. As shown in Fig. 2D, K2-AcrIIA4 fusions showed considerably reduced ability to repress dCasEV2.1-dependent luciferase activation in permissive temperature (14°C), however the remaining AcrIIA4 repression activity was reduced even further when incubated at restricted temperature (28°C), resulting in higher luminescence values indicative of partial degron functionality. To explore alternative regulatory options with wider dynamic range, we next analyzed N- and C-terminal fusions of the AcrIIA4 to the minimal auxin inducible degron (AID) mAID47^21^, again with or without the SV40 signal, as schematically depicted in Fig. 3A. AID was previously used for reprogramming *Arabidopsis thaliana* development when coupled to dCas9-based repressors^22^, as well as for inducing protein depletion in other eukaryotic systems^21,23,24^. Results in Fig. 3B show that most of the mAID fusions assayed partially retained the anti-Cas9 activity of native AcrIIA4, with some configurations showing almost full activity as it was the case for the mAID N-terminal fusion without nuclear localization signal (mAID-AcrIIA4). Upon auxin treatment, all protein fusions showed a clear de-repression trend indicative of auxin-mediated degradation, however the increase in reporter activity was only statistically significant for the above-mentioned mAID-AcrIIA4 fusion (four-fold activation). This is still an incomplete de-repression when compared with the luminescence levels obtained when AcrIIA4 is not present, thus indicating that this degron fusion needs further optimization to reach its full potential. To further confirm the auxin-dependency of the observed de-repression, we conducted a dose-response analysis. As shown in Fig. 3C, a near lineal dependency with the concentration of hormone in the treatment medium was clearly recorded. Auxin treatment was most effective when applied simultaneously with agroinfiltration, progressively losing its effect thereafter (Fig. 3D), suggesting that effective post-translation regulation occurs preferentially before the formation of the AcrIIA4-dCas9 complex.

**Figure 3.**
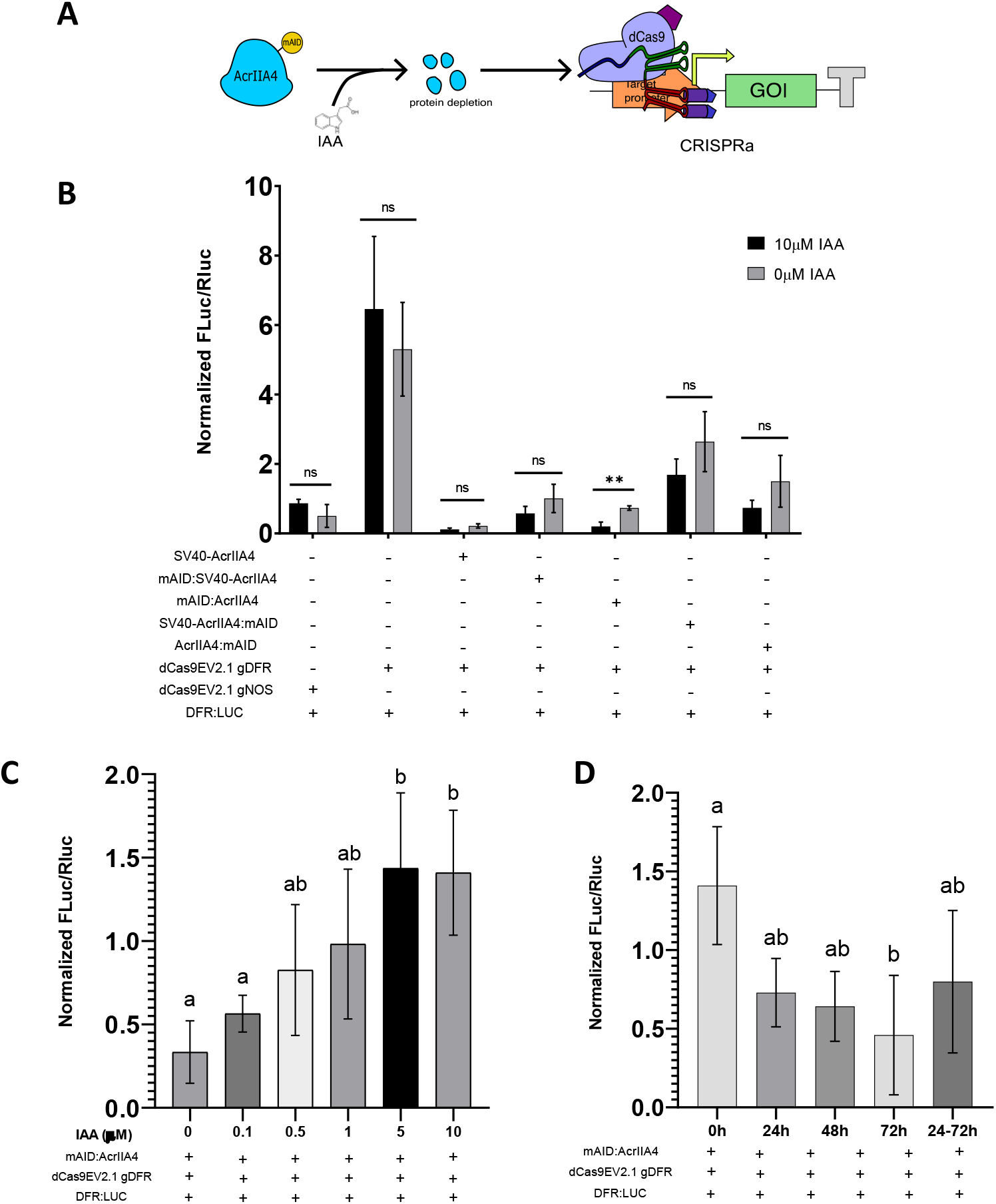
Auxin driven AcrIIA4 degradation confers tunable dCas9-based activation control in *N. benthamiana*. A) Schematic representation of IAA mediated AcrIIA4 protein depletion. B) Changes in transcriptional de-activation measured as normalized Fluc/Rluc ratios of *N. benthamiana* leaves expressing dCasEV2.1, the corresponding specific or unspecific gRNAs and four different versions of mAID-AcrIIA4 fusions in response to IAA treatments. C) Normalized Fluc/Rluc ratios of *N. benthamiana* leaves expressing dCasEV2.1, the corresponding specific or unspecific gRNAs and mAID:AcrIIA4 treated with different IAA concentrations or D) treated with 10μM IAA at different timepoints. SV40 is a nuclear localization signal. Error bars indicate SD (n=3). Statistical analyses were performed using One-way ANOVA (Tukey’s multiple comparisons test, P-Value ≤ 0.05) or unpaired t-Test for two samples comparisons (*P-value ≤ 0.05 and **P-value ≤ 0.005).

As a final application test, we analyzed the ability of AcrIIA4 to prevent dCasEV2.1-mediated gene activation of two endogenous *N. benthamiana* genes, Niben101Scf00305g05035 (*NbDFR*), encoding a dihydroflavonol reductase, and Niben101Scf00156g02004 (*NbAN2*), encoding a MYB factor involved in phenylpropanoid biosynthesis. Both genes were earlier shown to be efficiently activated with the dCasEV2.1 system^16^. *N. benthamiana* leaves were agroinfiltrated with the dCasEV2.1 system targeting *NbDFR* or *NbAN2* promoters with or without AcrIIA4. At 4 dpi leaf samples were collected and qRT-PCR assays were performed for each gene. To estimate gene expression levels, agroinfiltrated *NbDFR* samples were used as negative control for *NbAN2* samples and vice versa. As shown in Figure 4, the infiltration of dCasEV2.1 targeting *NbDFR* or *NbAN2* promoter regions conferred strong activation reaching up to 400 and 30 fold respectively, although the absolute levels were highly dependent on the age of the leaf. Remarkably, co-infiltration of AcrIIA4 strongly repressed CRISPR-mediated activation in both genes, resulting in a significant reduction of the relative gene expression values in all samples assayed.

**Figure 4.**
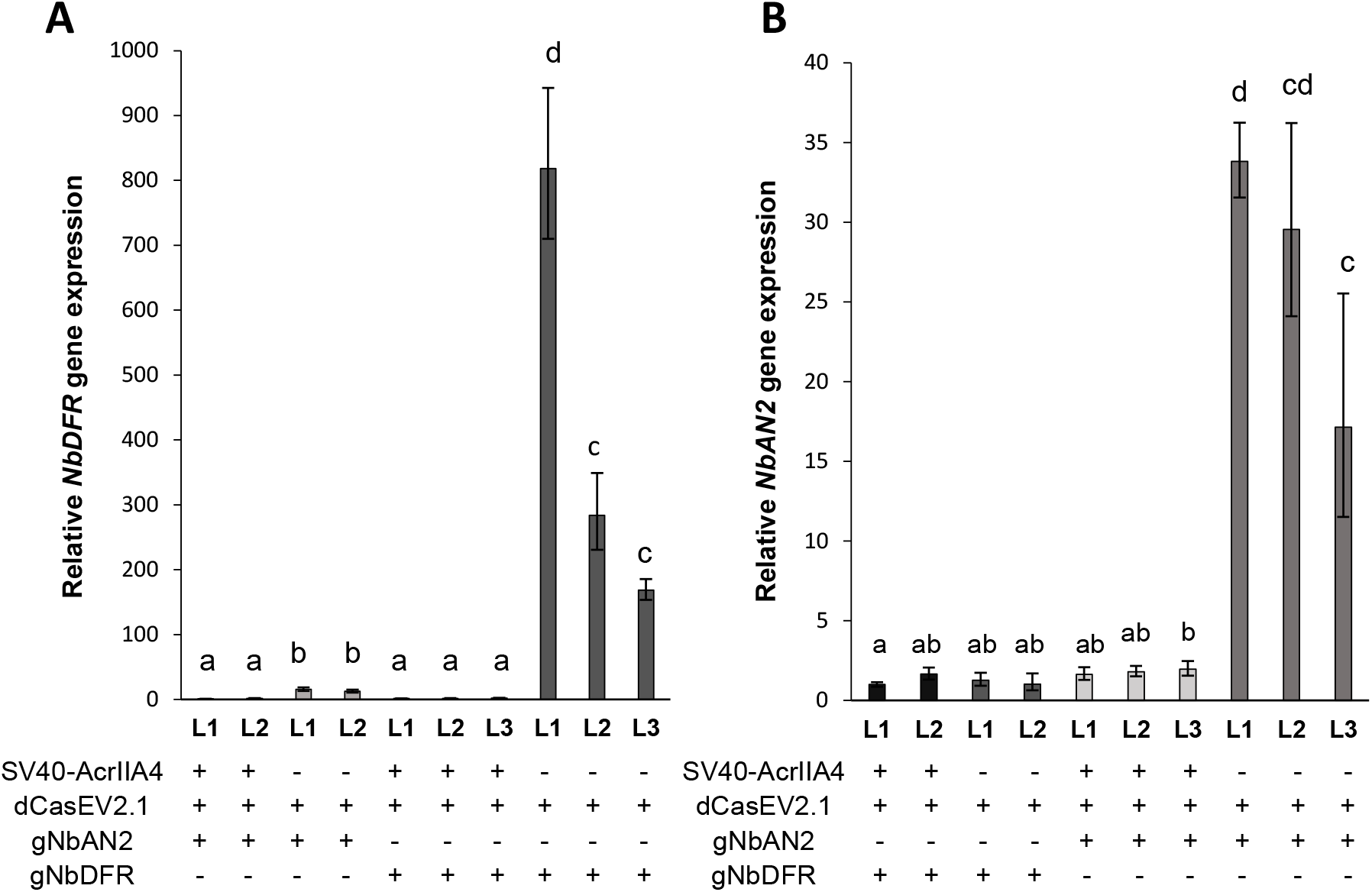
AcrIIA4 prevents dCas9-based activation of *NbDFR* and *NbAN2* in *N. benthamiana*. *N. benthamiana* leaves expressing dCas9EV2.1 and the corresponding specific or unspecific gRNAs for the activation of *NbDFR* or *NbAN2*, with or without SV40-AcrIIA4. L1, L2 and L3 indicate independent agroinfiltrated leaves. Error bars indicate SD (n=3, technical replicates). Statistical analyses were performed using One-way ANOVA (Tukey’s multiple comparisons test, P-Value ≤ 0.05).

In view of the results shown here, we conclude that AcrIIA4 is a potent tool for the spatial-temporal regulation of CRISPR/Cas9 activity in plants, Furthermore, in combination with programable transcriptional activation factors such as dCasEV2.1, AcrIIA4 will facilitate the customization of endogenous gene expression in plants.

## Materials and Methods

### GoldenBraid Cloning

DNA constructs used in this work were assembled using GoldenBraid^25^. AcrIIA4 protein sequence, as originally reported by Rauch et al.^26^ fused to SV40 NLS and mAID47 sequence, as reported by Brosh et al.^21^ (modified as N-tag or C-tag), were codon optimized for *N. benthamiana* using IDT Codon Optimization Tool (https://eu.idtdna.com/CodonOpt) and subsequently domesticated at https://gbcloning.upv.es/do/synthesis/. Additionally, a *N. benthamiana* codon optimized AcrIIA4 protein version without SV40-NLS, B3-B4 version, was domesticated using GoldenBraid GB Domesticator tool (https://gbcloning.upv.es/do/domestication/). Level 1 assemblies of transcriptional units from individual Level 0 parts were performed through Golden Gate-like multipartite BsaI reactions. All Level 0 and Level 1 assemblies were performed as previously reported ^19^. Level 0 assemblies were confirmed by restriction analysis and Sanger sequencing and Level 1 assemblies were verified by restriction analysis. An exhaustive list of all plasmids used in this work are listed in Supplementary Table 2.

### N. benthamiana transient expression

For transient expression assays, plasmids were transferred to *Agrobacterium tumefaciens* strain C58. Five to six weeks old *N. benthamiana* plants grown at 24°C and 16h (light)/ 20°C and 8h (darkness) conditions were used. Agroinfiltration was carried out as previously reported^27^. Briefly, overnight *A. tumefaciens* cultures were pelleted and resuspended in agroinfiltration solution (10 mM MES, pH 5.6, 10 mM MgCl_2_ and 200 μM acetosyringone) to an optical density of 0.1 at 600nm (OD_600_). Bacterial suspensions were incubated for 2h at room temperature on a horizontal rolling mixer, and then mixed for co-expression experiments, in which more than one GB element was used. Finally, agroinfiltrations were carried out through the abaxial surface of the three youngest fully expanded leaves of each plant with a 1ml needle-free syringe. For some experiments, agroinfiltrations were carried out with small modifications from the general procedure described above. For dose-response assays, SV40-AcrIIA4 TU (GB3344) was resuspended in agroinfiltration solution to an OD_600_ of 0.1 and subsequently diluted to 0.05, 0.01, 0.005, and 0.001 using a culture carrying an empty vector to maintain the final OD_600_ at 0.1. For the time-course assay, the strain carrying the SV40-AcrIIA4 TU (GB3344) was agroinfiltrated at OD_600_ of 0.05 at 0, 24, 48 and 72 h after infiltration of the dCasEV2.1 with the DFR gRNA (GB2513) and the pSlDFR reporter construct (GB1160). Detailed information of the experimental design can be found in the Supplementary Materials and Methods section.

### Luciferase activity and determination of relative transcriptional activity

Leaf samples were collected at 4 days post infiltration (dpi). For the determination of the FLuc/RLuc activity, one 0.8 cm diameter disc per agroinfiltrated leaf was excised. Leaf discs were frozen in liquid nitrogen and subsequently homogenized with 180 μl of Passive Lysis buffer, followed by 15 min of centrifugation at 14000 x g at 4°C. 10 μl of crude extract were mixed with 40 μl of LARII and Firefly luciferase (FLuc) activity was determined using a GloMax 96 microplate luminometer (Promega) with a 2-s delay and a 10-s measurement time. After the measurements, 40 μl of Stop&Glo Reagent were added per sample and Renilla luciferase (RLuc) activity was determined using the same protocol. Sample FLuc/RLuc ratios were calculated as the mean value of the three independent agroinfiltrated leaves. Relative transcriptional activities (RTAs) were calculated as the Fluc/Rluc ratios of the pSlDFR reporter in each sample normalized with the Fluc/Rluc ratios produced by a pNos reporter (GB1116) assayed in parallel and expressed in relative promoter units (rpu)^19^.

### Determination of Cas9-mediated editing activity

Genomic DNA was extracted from leaf samples 5 dpi following the CTAB protocol^28^. The DNA was used as template for PCR amplification of the targeted sites with primers listed on Supplementary Table 3 and using MyTaq™ DNA Polymerase (Bioline). Subsequently, PCR products were analyzed in 1% agarose gel electrophoresis, purified with the ExoSAP-IT™ PCR Product Cleanup Reagent (Applied Biosystems™) following manufacturer instructions and Sanger-sequenced. Finally, sequencing results were analyzed using Synthego CRISPR Performance Analysis (https://ice.synthego.com/#/) to determine the ICE score.

### RNA isolation and reverse transcription-quantitative PCR (RT-qPCR)

Leaf samples from infiltrated plants were harvested at 4 dpi and 100 mg of tissue were used for total RNA isolation using the Thermo Scientific™ GeneJET RNA Purification Kit. Total RNA was treated with Recombinant DNase I (RNase-free) (Takara) following manufacturer’s instructions. Aliquots of 1 μg of the treated RNA were used for cDNA synthesis with oligo dT using PrimeScript™ RT-PCR kit (Takara). cDNAs (0.4 μl) were used to determine the expression levels for each gene in triplicated 25 μl reactions with the SYBR® Premix Ex Taq (Takara) using the Applied biosystem 7500 Fast Real Time PCR system. *N. benthamiana* F-BOX gene was used as internal reference^29^. Calculations of each sample were carried out according the comparative ΔΔCT method^30^. Primers used for qRT-PCR reactions are listed in Supplementary Table 3.

## Acknowledgments

This work has been funded by PID2019-108203RB-100 Plan Nacional I+D, Spanish Ministry of Economy and Competitiveness. Vazquez-Vilar, M. is recipient of APOSTD/2020/096 (Generalitat Valenciana and Fondo Social Europeo post-doctoral grant). Selma, S. is recipient of a FPI fellowship (BIO2016‐78601‐R).

The authors declare no conflict of interest.

