## Supplementary Table 1, Supplementary Table 2, Supplementary Table 3, Supplementary Materials and Methods for "Strong and tunable anti-CRISPR/Cas9 activity of AcrIIA4 in plants"

#### Supplementary Information

**Supplementary Table 1.** Targeted genes and gRNA sequences of the editing experiment.

| Target gene | gRNA name | Protospacer sequence | Position |
|---|---|---|---|
| Niben101Scf04551g02001.1 (*NbXT2*) | gXT2 | GCTCTGATTGCACAATGGAA | +1628 |
| Niben101Scf01950g02001.1 (*SPL1950*) | gSPL | ATGGACATAACAGGCGTCGA | +562 |
| Niben101Scf00366g0008.1 (*SPL9294*) | | | +553 |

**Supplementary Table 2.** List of plasmids used in this work. Sequence information is available at http://www.gbcloning.upv.es/ by entering the GB Number.

| GB Number | Name |
|---|---|
| GB0107 | pDGB1α2:SF |
| GB0108 | pDGB3α1:P35S:p19:TNOS |
| GB0639 | pDGB2α2:P35S:hCas9:TNOS |
| GB1116 | pDGB3α1_PNOS:Luciferase:TNOS-SF-P35S:Renilla:TNOS-P35S:p19:TNOS |
| GB1119 | pDGB3α1_P35S:Luciferase:TNOS-SF-P35S:Renilla:TNOS-P35S:p19:TNOS |
| GB1160 | pDGB1Ω1_SlDFR:Luc:TNos-SF-35S:Renilla:TNos-35S:P19:Tnos |
| GB1770 | pDGB3α1_U6-26:gXT2:psgRNA |
| GB2049 | pDGB3α1_U6-26-4gRNA Pnos-F6x2_35s-dCas9:EDLL-Tnos-U6-26-5gRNA Pnos scf F6x2 - 35s-Ms2:VPR-Tnos |
| GB2513 | pDGB3α1_dCas9EDLL-Ms2:VPR SF -gRNA DFR -150 2.1 |
| GB3333 | pUPD2_SV40-AcrIIA4 (B3-B4-B5) |
| GB3659 | pUPD2_SV40-AcrIIA4 (B3-B4) |
| GB3660 | pUPD2_AcrIIA4 (B3-B4) |
| GB3661 | pUPD2_AcrIIA4 (B3-B4-B5) |
| GB3663 | pUPD2_mAID (N-tag) B2 |
| GB3664 | pUPD2_mAID (C-tag) B5 |
| GB3344 | pDGB3α1:P35S:SV40-AcrIIA4:TNOS |
| GB3666 | pDGB3α1_P35S:mAID:SV40-AcrIIA4:TNOS |
| GB3667 | pDGB3α1_P35S:SV40-AcrIIA4:mAID:TNOS |
| GB3668 | pDGB3α1_P35S:mAID:AcrIIA4:TNOS |
| GB3669 | pDGB3α1:p35S:AcrIIA4:mAID:TNOS |
| GB3670 | pDGB3α1_P35S:K2:SV40-AcrIIA4:TNOS |
| GB3671 | pDGB3α1_P35S:K2:AcrIIA4:TNOS |
| GB3701 | pDGB3α1:U6-26:gSPL1.6:psgRNA |

**Supplementary Table 3.** List of primers used in this work.

| Gene identifier | Primer Sequence (5'→ 3') |
|---|---|
| NbXT2 | TGCACGGTTGTCCGAGTTTG |
| | TTACTTGTGAATTGCTCTCTGGT |
| SPL1950 | AAATGTTCAATCCCTGGACGAC |
| | CAAAACGGACAGCGACATGGT |
| SPL9294 | GCCCAGGTTTGAATGCATTAGGG |
| | GAAAACGGACAACGACATGGT |
| NbDFR | TTCATCTGCGCATCCCATCA |
| | TCCCTACTGAGTTTAAAGGTATCGA |
| NbAN2 | GGAAAAGTTGCAGACTGAGGTG |
| | ACCCGCAATAAGTGACCATCTG |
| NbF-box | TTGGAAACTCTCTCCCCACTTG |
| | GCTCATTGTTGGATGGGTACCT |

#### Supplementary Materials and Methods

The tables provided below show the Agrobacterium cultures co-infiltrated in the same mix for each experiment.

*Directed gene editing assay (Figure 1B)*

| Construct | Plant 1 | Plant 2 | Plant 3 | Plant 4 |
|---|---|---|---|---|
| 35S:SV40-AcrIIA4:TNos (GB3344) | + | - | + | - |
| 35s:hCas9:tNos (GB0639) | + | + | + | + |
| U6-26:gXT2:psgRNA (GB1770) | + | + | - | - |
| U6-26:gSPL1.6:psgRNA (GB3701) | - | - | + | + |
| pSF (GB0107) | - | + | - | + |
| 35s:P19:TNos (GB0108) | + | + | + | + |

*Luciferase reporter activation assay (Figure 1C)*

| Construct | Plant 1 | Plant 2 | Plant 3 | Plant 4 | Plant 5 |
|---|---|---|---|---|---|
| 35S:SV40-AcrIIA4:Tnos (GB3344) | + | - | - | - | - |
| SlDFR:Luc:TNos-SF-35S:Renilla:TNos-35S:P19:Tnos (GB1160) | + | + | + | - | - |
| dCas9EDLL-Ms2:VPR SF -gRNA DFR -150 2.1 (GB2513) | + | + | - | - | - |
| U6-26-4gRNA Pnos-F6x2_35s-dCas9:EDLL-Tnos-U6-26-5gRNA Pnos scf F6x2 - 35s-Ms2:VPR-Tnos (GB2049) | - | - | + | - | - |
| pSF (GB0107) | - | + | + | - | - |
| PNos:Luciferase:TNos-SF-35S:Renilla:TNos-35S:P19:TNos-SF (GB1116) | - | - | - | + | - |

*Dose-response assay (Figure 2A)*

| Construct | Plant 1 | Plant 2 | Plant 3 | Plant 4 | Plant 5 | Plant 6 | Plant 7 | Plant 8 | Plant 9 | Plant 10 | Plant 11 |
|---|---|---|---|---|---|---|---|---|---|---|---|
| 35S:SV40-AcrIIA4:Tnos (GB3344) OD 0.1 | + | - | - | - | - | - | - | - | - | - | - |
| 35S:SV40-AcrIIA4:Tnos (GB3344) OD 0.05 | - | + | - | - | - | - | - | - | - | - | - |
| 35S:SV40-AcrIIA4:Tnos (GB3344) OD 0.01 | - | - | + | - | - | - | - | - | - | - | - |
| 35S:SV40-AcrIIA4:Tnos (GB3344) OD 0.005 | - | - | - | + | - | - | - | - | - | - | - |
| 35S:SV40-AcrIIA4:Tnos (GB3344) OD 0.001 | - | - | - | - | + | - | - | - | - | - | - |
| 35S:SV40-AcrIIA4:Tnos (GB3344) OD 0.0005 | - | - | - | - | - | + | - | - | - | - | - |
| SlDFR:Luc:TNos-SF-35S:Renilla:TNos-35S:P19:Tnos (GB1160) | + | + | + | + | + | + | + | + | - | - | - |
| dCas9EDLL-Ms2:VPR SF -gRNA DFR -150 2.1 (GB2513) | + | + | + | + | + | + | + | - | - | - | - |
| U6-26-4gRNA Pnos-F6x2_35s-dCas9:EDLL-Tnos-U6-26-5gRNA Pnos scf F6x2 - 35s-Ms2:VPR-Tnos (GB2049) | - | - | - | - | - | - | - | + | - | - | - |
| pSF (GB0107) | - | - | - | - | - | - | + | + | - | - | - |
| PNos:Luciferase:TNos-SF-35S:Renilla:TNos-35S:P19:TNos-SF (GB1116) | - | - | - | - | - | - | - | - | + | - | - |

#### *Time-course assay (Figure 2B)*

| **Construct** | **Plant 1 (day 1)** | **Plant 2 (day 2)** | **Plant 3 (day 3)** | **Plant 4 (day4)** | **Plant 5** | **Plant 6** | **Plant 7** | **Plant 8** |
|---|---|---|---|---|---|---|---|---|
| 35S:SV40-AcrIIA4:Tnos (GB3344) | + | + | + | + | - | - | - | - |
| SlDFR:Luc:TNos-SF-35S:Renilla:TNos-35S:P19:Tnos (GB1160) | + | + | + | + | + | + | - | - |
| dCas9EDLL-Ms2:VPR SF -gRNA DFR -150 2.1 (GB2513) | + | + | + | + | + | - | - | - |
| U6-26-4gRNA Pnos-F6x2_35s-dCas9:EDLL-Tnos-U6-26-5gRNA Pnos scf F6x2 - 35s-Ms2:VPR-Tnos (GB2049) | - | - | - | - | - | + | - | - |
| pSF (GB0107) | - | - | - | - | + | + | - | - |
| PNos:Luciferase:TNos-SF-35S:Renilla:TNos-35S:P19:TNos-SF (GB1116) | - | - | - | - | - | - | + | - |

#### *Temperature Dependent Degron assay (Figure 2D)*

| | **14°C** | | | | **28°C** | | | |
|---|---|---|---|---|---|---|---|---|
| **Construct** | **Plant 1** | **Plant 2** | **Plant 3** | **Plant 4** | **Plant 5** | **Plant 6** | **Plant 7** | **Plant 8** |
| 35S:K2:SV40-AcrIIA4:Tnos (GB3670) OD0.05 | **+** | **-** | **-** | **-** | **+** | **-** | **-** | **-** |
| 35S:K2:AcrIIA4:Tnos (GB3671) OD0.05 | - | + | + | - | - | + | + | - |
| SlDFR:Luc:TNos-SF-35S:Renilla:TNos-35S:P19:Tnos (GB1160) | + | + | + | - | + | + | + | - |
| dCas9EDLL-Ms2:VPR SF -gRNA DFR -150 2.1 (GB2513) | + | + | - | - | + | + | - | - |
| U6-26-4gRNA Pnos-F6x2_35s-dCas9:EDLL-Tnos-U6-26-5gRNA Pnos scf F6x2 - 35s-Ms2:VPR-Tnos (GB2049) | - | - | + | - | - | - | + | - |
| pSF (GB0040) | + | + | + | - | + | + | + | - |
| PNos:Luciferase:TNos-SF-35S:Renilla:TNos-35S:P19:TNos-SF (GB1116) | - | - | - | + | - | - | - | + |

### *Auxin Inducible Degron assay (Figure 3B)*

| | *10µM IAA* | | | | | | | | *0µM IAA* | | | | | | | | |
|---|---|---|---|---|---|---|---|---|---|---|---|---|---|---|---|---|---|
| **Construct** | ***Plant 1*** | ***Plant 2*** | ***Plant 3*** | ***Plant 4*** | ***Plant 5*** | ***Plant 6*** | ***Plant 7*** | ***Plant 8*** | ***Plant 9*** | ***Plant 10*** | ***Plant 11*** | ***Plant 12*** | ***Plant 13*** | ***Plant 14*** | ***Plant 15*** | ***Plant 16*** | ***Plant 17*** |
| 35S:SV40-AcrIIA4:Tnos (GB3344) OD0.05 | - | - | - | - | + | - | - | - | - | - | - | - | + | - | - | - | - |
| 35S:mAID:SV40-AcrIIA4:Tnos (GB3666) OD0.05 | + | - | | - | | - | - | - | + | - | | - | | - | - | - | - |
| 35S:SV40-AcrIIA4:mAID:Tnos (GB3667) OD0.05 | - | + | - | - | - | - | - | - | - | + | - | - | - | - | - | - | - |
| 35S:mAID:AcrIIA4:Tnos (GB3668) OD0.05 | - | - | + | - | - | - | - | - | - | - | + | - | - | - | - | - | - |
| 35S:AcrIIA4:mAID:Tnos (GB3669) OD0.05 | - | - | - | + | - | - | - | - | - | - | - | + | - | - | - | - | - |
| SlDFR:Luc:TNos-SF-35S:Renilla:TNos-35S:P19:Tnos (GB1160) | + | + | + | + | + | + | + | - | + | + | + | + | + | + | + | - | - |
| dCas9EDLL-Ms2:VPR SF -gRNA DFR -150 2.1 (GB2513) | + | + | + | + | + | + | - | - | + | + | + | + | + | + | - | - | - |
| U6-26-4gRNA Pnos-F6x2_35s-dCas9:EDLL-Tnos-U6-26-5gRNA Pnos scf F6x2 - 35s-Ms2:VPR-Tnos (GB2049) | - | - | - | - | - | - | + | - | - | - | - | - | - | - | + | - | - |
| pSF (GB0107) | - | + | - | - | - | + | + | - | - | + | - | - | - | + | + | - | - |
| PNos:Luciferase:TNos-SF-35S:Renilla:TNos-35S:P19:TNos-SF (GB1116) | - | - | - | - | - | - | - | + | - | - | - | - | - | - | - | + | - |

### *Auxin Inducible Degron dose-response assay (Figure 3C)*

| **Construct** | ***Plant 1 (5 µM)*** | ***Plant 2 (1 µM)*** | ***Plant 3 (0.5µM)*** | ***Plant 4 (0.1µM)*** | ***Plant 5*** | ***Plant 6*** | ***Plant 7*** | ***Plant 8*** |
|---|---|---|---|---|---|---|---|---|
| 35S:mAID:AcrIIA4:Tnos (GB3668) OD0.05 | + | + | + | + | - | - | - | - |
| SlDFR:Luc:TNos-SF-35S:Renilla:TNos-35S:P19:Tnos (GB1160) | + | + | + | + | + | + | - | - |
| dCas9EDLL-Ms2:VPR SF -gRNA DFR -150 2.1 (GB2513) | + | + | + | + | + | - | - | - |
| U6-26-4gRNA Pnos-F6x2_35s-dCas9:EDLL-Tnos-U6-26-5gRNA Pnos scf F6x2 - 35s-Ms2:VPR-Tnos (GB2049) | - | - | - | - | - | + | - | - |
| pSF (GB0107) | + | + | + | + | + | + | - | - |
| PNos:Luciferase:TNos-SF-35S:Renilla:TNos-35S:P19:TNos-SF (GB1116) | - | - | - | - | - | - | + | - |

*Auxin Inducible Degron time-course assay (Figure 3D)*

| ***Construct*** | ***Plant 1 (0h)*** | ***Plant 2 (24h)*** | ***Plant 3 (48h)*** | ***Plant 4 (72h)*** | ***Plant 5 (24-72h)*** | ***Plant 6*** | ***Plant 7*** | ***Plant 8*** | ***Plant 9*** |
|---|---|---|---|---|---|---|---|---|---|
| 35S:mAID:AcrIIA4:Tnos (GB3668) OD0.05 | + | + | + | + | + | - | - | - | - |
| SlDFR:Luc:TNos-SF-35S:Renilla:TNos-35S:P19:Tnos (GB1160) | + | + | + | + | + | + | + | - | - |
| dCas9EDLL-Ms2:VPR SF -gRNA DFR -150 2.1 (GB2513) | + | + | + | + | + | + | - | - | - |
| U6-26-4gRNA Pnos-F6x2_35s-dCas9:EDLL-Tnos-U6-26-5gRNA Pnos scf F6x2 - 35s-Ms2:VPR-Tnos (GB2049) | - | - | - | - | - | - | + | - | - |
| pSF (GB0107) | + | + | + | + | + | + | + | - | - |
| PNos:Luciferase:TNos-SF-35S:Renilla:TNos-35S:P19:TNos-SF (GB1116) | - | - | - | - | - | - | - | + | - |

*Endogenous genes activation assay (Figure 4)*

| ***Construct*** | ***Plant 1*** | ***Plant 2*** | ***Plant 3*** | ***Plant 4*** |
|---|---|---|---|---|
| 35S:SV40-AcrIIA4:TNos (GB3344) | + | - | + | - |
| 35s-Ms2:VPR-Tnos-35s-dCas9:EDLL-Tnos (GB2085) | + | + | + | + |
| U6-26-sgRNANbDFR -85,-198,-268 F6x2 Multiplex (GB2170) | + | + | - | - |
| U6-26-sgRNANbAN2 -103,-175,-196 F6x2 (GB2171) | - | - | + | + |
| pSF (GB0040) | - | + | - | + |
| 35s:P19:TNos (GB0108) | + | + | + | + |
